# Modulation masking and fine structure shape neural envelope coding to predict speech intelligibility across diverse listening conditions

**DOI:** 10.1101/2021.03.26.437273

**Authors:** Vibha Viswanathan, Hari M. Bharadwaj, Barbara G. Shinn-Cunningham, Michael G. Heinz

## Abstract

A fundamental question in the neuroscience of everyday communication is how scene acoustics shape the neural processing of attended speech sounds and in turn impact speech intelligibility. While it is well known that the temporal envelopes in target speech are important for intelligibility, how the neural encoding of target-speech envelopes is influenced by background sounds or other acoustic features of the scene is unknown. Here, we combine human electroencephalography with simultaneous intelligibility measurements to address this key gap. We find that the neural envelope-domain signal-to-noise ratio in target-speech encoding, which is shaped by masker modulations, predicts intelligibility over a range of strategically chosen realistic listening conditions unseen by the predictive model. This provides neurophysiological evidence for modulation masking. Moreover, using high-resolution vocoding to carefully control peripheral envelopes, we show that target-envelope coding fidelity in the brain depends not only on envelopes conveyed by the cochlea, but also on the temporal fine structure (TFS), which supports scene segregation. Our results are consistent with the notion that temporal coherence of sound elements across envelopes and/or TFS influences scene analysis and attentive selection of a target sound. Our findings also inform speech-intelligibility models and technologies attempting to improve real-world speech communication.

## 1 Introduction

A fundamental question in sensory neuroscience is how our brains parse complex scenes to organize the barrage of sensory information into perceptually coherent objects and streams. Low-level regularities in stimulus features, such as proximity and continuity of boundaries/textures in vision (Gray, 1999) or rhythmicity, pitch, and harmonicity in audition (Darwin, 1997), can promote perceptual binding and scene segregation. Speech perception in complex environments is a prominent example where such feature-based scene analysis is critical for everyday communication (Cherry, 1953), yet the neurophysiological mechanisms supporting this process are poorly understood. Elucidating the mechanisms underlying speech intelligibility is important for both clinical applications and audio technologies, such as optimizations for cochlear-implant and hearing-aid signal processing, clinical diagnostics and individualized interventions for speech-in-noise communication problems, and speech denoising algorithms (e.g., in cell phones).

Any acoustic signal can be decomposed into a slowly varying temporal modulation, or envelope, and a rapidly varying temporal fine structure (TFS) (Hilbert, 1906). In the auditory system, the cochlea decomposes broadband inputs into a tonotopic representation, where each channel encodes the signal content in a relatively narrow band of frequencies around a different center frequency. The envelope and TFS information in each channel is then encoded through the activity of neurons in the ascending auditory pathway (Johnson, 1980; Joris and Yin, 1992). Psychophysical studies suggest that envelopes convey important information about speech content (Shannon et al., 1995; Smith et al., 2002; Elliott and Theunissen, 2009), whereas TFS is important for our perception of attributes such as pitch and location (Smith et al., 2002).

The temporal coherence theory of scene analysis (Elhilali et al., 2009; Gray, 1999) suggests that neural assemblies that fire coherently (driven by envelopes, TFS, or both) support perceptual grouping of sound elements across distinct frequency channels, which can aid source segregation (Schooneveldt and Moore, 1987). This may also explain how masker elements that are temporally coherent with target speech, but are in a different channel from the target can perceptually interfere (Apoux and Bacon, 2008). Accordingly, the temporal coherence theory makes important predictions about how the envelopes and TFS of sources in a scene affect scene analysis and thus how they should influence the neural representation and intelligibility of target speech. However, these predictions have not been evaluated in neurophysiological experiments for realistic listening conditions that capture the complexity of everyday “cocktail-party” environments.

A parallel psychoacoustic literature suggests that modulation masking (i.e., the internal representation of temporal modulations in the target relative to those from the background, which contains inherent distracting fluctuations) may be a key contributor to speech understanding in noise (Bacon and Grantham, 1989; Stone and Moore, 2014). Accordingly, while classic speech-intelligibility models emphasized audibility in different frequency bands (ANSI, 1969, 1997), current models that emphasize envelope coding (Steeneken and Houtgast, 1980) and modulation masking (Dubbelboer and Houtgast, 2008; Relaño-Iborra et al., 2016) have been successful in predicting performance in many realistic conditions. However, the core notion that modulation masking is important has not been validated neurophysiologically. With the exception of current speech-intelligibility models that restrict modulation masking effects to within a carrier frequency channel (Jørgensen et al., 2013; Relaño-Iborra et al., 2016), the literature on modulation masking largely does not distinguish between cross-channel interference and within-channel masking. In this sense, the theory of modulation masking is consistent with the temporal coherence theory. However, modulation masking does not consider the role of TFS, despite the consistent finding that cues conveyed by TFS (e.g., pitch) (Smith et al., 2002) critically support object formation, perceptual scene segregation, and selective attention (Darwin, 1997; Shinn-Cunningham, 2008). Indeed, temporal coherence across low-frequency TFS and high-frequency pitch envelopes may significantly improve speech intelligibility in noise, compared to having either cue alone (Oxenham and Simonson, 2009). While some psychophysical studies have explored the relative roles of envelope and fine-structure cues for speech intelligibility in noise (Qin and Oxenham, 2003; Lorenzi et al., 2006; Swaminathan and Heinz, 2012), few neurophysiological studies have investigated how these cues work together during selective listening.

In the present study, we bridge these gaps by measuring electroencephalography (EEG) simultaneously with intelligibility for target speech over a range of strategically chosen realistic listening conditions. The EEG measured is the response evoked by ongoing stimulus fluctuations when attending to the target speech. We hypothesized that the neural tracking of target modulations, as quantified from EEG, will depend strongly on the modulation content of the masker, in line with the temporal coherence theory and the notion of modulation masking. Furthermore, we hypothesized that the availability (or lack thereof) of TFS will also impact this neural target-envelope coding, in line with the role of TFS in providing cues to facilitate scene analysis and attention. Finally, we hypothesized that the net neural target-envelope coding shaped by these factors [i.e., the neural signal-to-noise ratio (SNR) in the envelope domain] will predict (in a quantitative, statistical sense) speech intelligibility in conditions unseen by the predictive model. Our neurophysiological results provide evidence for all of the above hypotheses. The present study thus goes beyond comparing individual outcomes to neural measures in a particular condition (e.g., Ding and Simon, 2013; Bharadwaj et al., 2015), to elucidate what aspects of the scene acoustics and neural processing predict intelligibility across diverse real-world conditions.

## 2 Materials and Methods

### 2.1 Stimulus generation

Seven hundred Harvard/Institute of Electrical and Electronics Engineers (IEEE) sentences (Rothauser, 1969) spoken in a female voice and recorded as part of the PN/NC corpus (McCloy et al., 2013) were chosen for the study. The Harvard/IEEE lists have relatively low semantic context compared to other commonly used speech material (Boothroyd and Nittrouer, 1988; Rabinowitz et al., 1992; Grant and Seitz, 2000). Stimuli were created for eight different experimental conditions as described below:

1-3. **Speech in speech-shaped stationary noise (SiSSN)**: Speech was added to spectrally matched stationary gaussian noise, i.e., speech-shaped stationary noise, at SNRs of -2, - 5, and -8 dB. The long-term spectra of the target speech sentences and that of stationary noise were adjusted to match the average (across instances) long-term spectrum of four-talker babble. A different realization of stationary noise was used for each SiSSN stimulus.
4-5. **Speech in babble (SiB)**: Speech was added to spectrally matched four-talker babble at SNRs of 4 and -2 dB. The long-term spectra of the target speech sentences were adjusted to match the average (across instances) long-term spectrum of four-talker babble. In creating each SiB stimulus, a babble sample was randomly selected from a list comprising 72 different four-talker babble maskers obtained from the QuickSIN corpus (Killion et al., 2004).
6. **Speech in babble with reverberation (SiB reverb)**: SiB at 6 dB SNR was subjected to reverberation simulating St. Albans Cathedral in England [by convolution with a binaural impulse response; see Gorzel et al. (2010)]. The reverberation time (T60) was 2.4 s.
7. **Vocoded speech in babble (SiB vocoded)**: SiB at 4 dB SNR was subjected to 64-channel envelope vocoding, which left the peripheral envelopes and place coding intact, while replacing the TFS with a noise carrier in accordance with the procedure described in Qin and Oxenham (2003). The 64 frequency channels were contiguous with their center frequencies equally spaced on an ERB-number scale (Glasberg and Moore, 1990) between 80 and 6000 Hz. To verify that the vocoding procedure did not significantly change envelopes at the cochlear level, we extracted the envelopes at the output of 128 filters (using a similar procedure as in the actual vocoding process) both before and after vocoding for 50 different SiB stimuli. Note that the use of 128 filters allowed us to compare envelopes at both on-band filters (i.e., filters whose center frequencies matched with the sub-bands of the vocoder) and off-band filters (i.e., filters whose center frequencies were halfway between adjacent vocoder sub-bands on the ERB-number scale). The average correlation coefficient between envelopes before and after vocoding (across the 50 SiB stimuli and the 128 cochlear filters and after adjusting for any vocoder group delays) is about 0.9. This suggests that our vocoding procedure leaves the cochlear-level envelopes largely intact. Indeed, as illustrated in Figure 1, our 64-channel vocoding procedure better preserves the within-band envelopes than the lower-resolution procedures of Ding et al. (2014). Although Ding et al. (2014) suggested that TFS matters for neural envelope tracking, their methods using four- or eight-channel vocoding do not preserve peripheral envelopes within individual cochlear bands. Consequently, a purely envelope-based explanation of their findings cannot be ruled out.

**Figure 1.**
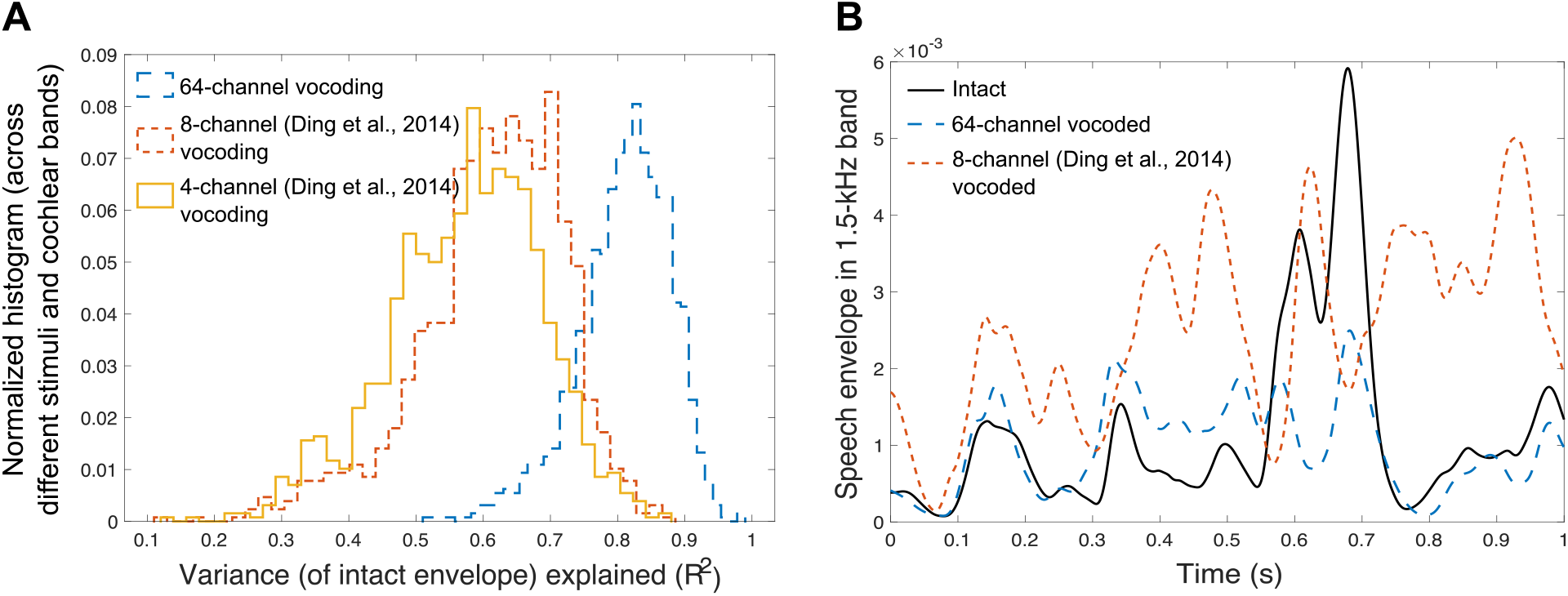
Illustration of the effect of 64-channel vocoding versus the lower-resolution procedures of Ding et al. (2014) on envelopes within individual cochlear bands. **Panel A** shows a histogram of the group-delay-adjusted squared normalized-correlation (i.e., variance explained) between the envelope in intact speech in babble (SiB) and 64-channel vocoded SiB, which is used in this present study (i), and the eight-channel (ii) and four-channel (iii) vocoding of Ding et al. (2014) vocoded SiB. The histograms are across different speech sentences and 128 different cochlear bands equally spaced on an ERB-number scale (Glasberg and Moore, 1990) from 80-6000 Hz. The 64-channel vocoding clearly better preserves the within-band envelopes than either the eight- or four-channel procedures of Ding et al. (2014) in that the 64-channel procedure captures an additional variance of more than 20%. This disruption of within-band envelopes using their technique was observed despite replicating their result of 0.99 correlation for the band-summed envelope (i.e., the basis for their conclusion that their vocoding preserved speech envelopes). **Panel B** shows an example envelope derived from SiB for the 1.5-kHz speech band for intact SiB, our 64-channel vocoding, and the better-resolution, eight-channel vocoding from Ding et al. (2014), to visualize how our procedure yields band-specific envelopes that more closely match those of intact SiB.
8. **Speech in babble with ideal time-frequency segregation (SiB ITFS)**: SiB at -6 dB SNR was subjected to 64-channel ITFS, a non-linear denoising procedure that forms the basis of many machine-learning denoising strategies (Wang and Chen, 2018). This was performed over a frequency range of 80–8000 Hz, mirroring the procedure in Brungart et al. (2006). A local SNR criterion of 0 dB was used in the ITFS procedure.

Prior to the full study, a behavioral pilot study (with five subjects who did not participate in the actual EEG experiment) was used to determine the SNRs for the different experimental conditions. The SNRs for the three SiSSN conditions were chosen to yield intelligibility values of roughly 25%, 50% and 75%, respectively, to span the full range of intelligibility. The SNRs for the other conditions were chosen such that the intelligibility scores were between (but did not include) 0% and 100% and were different across the different conditions.

Table 1 lists the different stimulus conditions along with the rationale for including them in our study.

**Table 1.** Rationale for the different stimulus conditions included in this study. Collectively, the different listening conditions represent a diversity of scene acoustics, including important examples in our environment and clinical applications. Moreover, they span maskers with different modulation statistics (Jørgensen et al., 2013; Rosen et al., 2013) and stimuli with intact and degraded TFS, which allowed us to rigorously test our hypotheses. Note that the SNR levels were chosen to span the full range of intelligibility without floor or ceiling effects.

| No. | Stimulus condition | Rationale for inclusion in study |
| --- | --- | --- |
| 1-3 | Speech in Speech-shaped Stationary Noise (SiSSN) at SNRs of -2, -5, and -8 dB | Widely used in the literature; used for calibration of prediction model |
| 4-5 | Speech in Babble (SiB) at SNRs of 4 and -2 dB | Simulates ecologically relevant cocktail-party listening; has different masker modulation statistics from SiSSN |
| 6 | SiB at 6 dB SNR subjected to reverberation (T60 = 2.4 s) | Reverberation is ubiquitous in everyday listening environments (e.g., rooms and stairwells); linearly distorts temporal information |
| 7 | SiB at 4 dB SNR subjected to 64-channel envelope vocoding | Used to investigate the role of TFS in target-speech coding and intelligibility |
| 8 | SiB at -6 dB SNR subjected to 64-channel ideal time-frequency segregation (ITFS) | ITFS is a precursor to deep-learning-based denoising algorithms that are increasingly used in many audio processing applications, including hearing aids (Wang and Chen, 2018); nonlinear distortion |

### 2.2 Participants

Data were collected from 12 human subjects (four male), aged 19–31 years, recruited from the Purdue University community. All subjects were native speakers of American English, had pure-tone hearing thresholds better than 20 dB hearing level in both ears at standard audiometric frequencies between 250 Hz and 8 kHz, and reported no neurological disorders. All subjects also had distortion-product and click-evoked otoacoustic emissions (DPOAE and CEOAE) within the normal range of published values for individuals with normal hearing (Gorga et al., 1993), as well as normal tympanograms. Subjects provided informed consent in accordance with protocols established at Purdue University. Data were collected from each subject over the course of one or two visits (with a total visit time of *∼*5 h).

### 2.3 Experimental design

Each subject performed seven blocks of speech intelligibility testing, with 100 trials per block, and with a distinct target sentence in each trial. Subjects had a 5–10-min break between successive blocks. Different but overlapping subsets of experimental conditions were randomly assigned to each subject, such that at least 700 trials for each experimental condition were collected across the subject cohort. This design avoided confounding individual-subject effects with experimental-condition effects. The different experimental conditions were intermingled within each block.

Subjects were instructed that in each trial they would be listening for a woman’s voice speaking a sentence and that at the end of the trial they would have to verbally repeat the sentence back to the experimenter sitting beside them in a sound-treated booth. They were told that it would be the same woman’s voice every time but that the type and level of background noise/distortion would change from trial to trial. They were also instructed that in each trial, the noise would start first and the target woman’s voice *∼*1 s later. They were encouraged to guess as many words as they could if they heard a sentence only partially.

Stimuli were presented to subjects diotically in all conditions except the reverberation condition, in which stimuli were generated with ear-specific impulse responses as described previously. Thirty-two-channel EEG was measured while subjects performed the behavioral task. The target speech sentences were presented at a sound level of 72 dB sound pressure level (SPL), while the level of the background was varied according to the stimulus SNR.

At the beginning of each trial, subjects were presented with a visual cue that read “stay still and listen now” in red font. The audio stimulus started playing 1 s after the visual cue was presented. In every stimulus presentation, the background noise started first and continued for the entire duration of the trial, while the target speech started 1.25 s after the background started. This was done to help cue the subjects’ attention to the stimulus before the target sentence was played. The target was at least 2.5 s long. After the target sentence ended, the background noise continued for a short amount of time that varied randomly from trial to trial. This was done to reduce EEG contamination from movement artifacts and motor-planning signals. Two hundred ms after the noise ended, subjects were presented with a different visual cue that read “repeat now” in green font, cueing them to report the target sentence to the experimenter. Intelligibility was scored on five pre-determined keywords (which excluded articles and prepositions) for each sentence. For each experimental condition, an overall intelligibility score was obtained by averaging the percentage of key words correct (for a sentence) over all sentences used in that condition and across subjects.

Subjects performed a short training demo task before the actual EEG experiment. The demo spanned the same set of listening conditions and used the same woman’s voice as the actual experiment but contained a different set of Harvard/IEEE target sentences, not used in the main experiment. All 12 subjects scored more than 70% on the easiest condition and got at least some words correct (*>* 0%) on the hardest condition. All were able to stay still during the presentation of the sentences and respond on cue. This ensured that in the actual experiment, intelligibility scores showed minimal ceiling or floor effects and that movement artifacts were minimal, providing clean EEG recordings.

### 2.4 Hardware

A personal desktop computer controlled all aspects of the experiment, including triggering sound delivery and storing data. Special-purpose sound-control hardware (System 3 real-time signal processing system, including digital-to-analog conversion and amplification; Tucker Davis Technologies, Alachua, FL) presented audio through insert earphones (ER-2; Etymotic, Elk Grove Village, IL) coupled to foam ear tips. The earphones were custom shielded by wrapping the transducers in layers of magnetic shielding tape made from an amorphous cobalt alloy (MCF5; YSHIELD GmbH & Co., Ruhstorf, Germany) and then placing them in 3-mm-thick aluminum enclosures to attenuate electromagnetic interference. The signal cables driving the transducers were shielded with braided metallic Techflex (Techflex, Sparta, NJ). All shielding layers were grounded to the chassis of the digital-to-analog (D/A) converter. The absence of measurable electromagnetic artifact was verified by running intense click stimuli through the transducers with the transducers positioned in the same location relative to the EEG cap as actual measurements but with foam tips left outside the ear. All audio signals were digitized at a sampling rate of 48.828 kHz. The EEG signals were recorded at a sampling rate of 4.096 kHz using a BioSemi (Amsterdam, The Netherlands) ActiveTwo system. Recordings were done with 32 cephalic electrodes and two additional earlobe electrodes.

### 2.5 Data preprocessing

Of the 12 subjects who participated, one subject could not complete the task because they were sleepy, and another subject was unable to return for their second visit to complete the study. Data from these two subjects were excluded from the study. The EEG signals of the remaining ten subjects were re-referenced to the average of the two earlobe reference electrodes. The Signal Space Projection method was used to construct spatial filters to remove eye blink and saccade artifacts (Uusitalo and Ilmoniemi, 1997). The broadband EEG was then bandpass filtered between 1 and 400 Hz for further analysis. Data from three completely new subjects (who were not among the 12 who participated in the main experiment) showed that responses from the auditory cortex and brainstem are strongest in EEG channel FCz (see Fig. 2B); thus, we used FCz to derive all results presented in this report.

**Figure 2.**
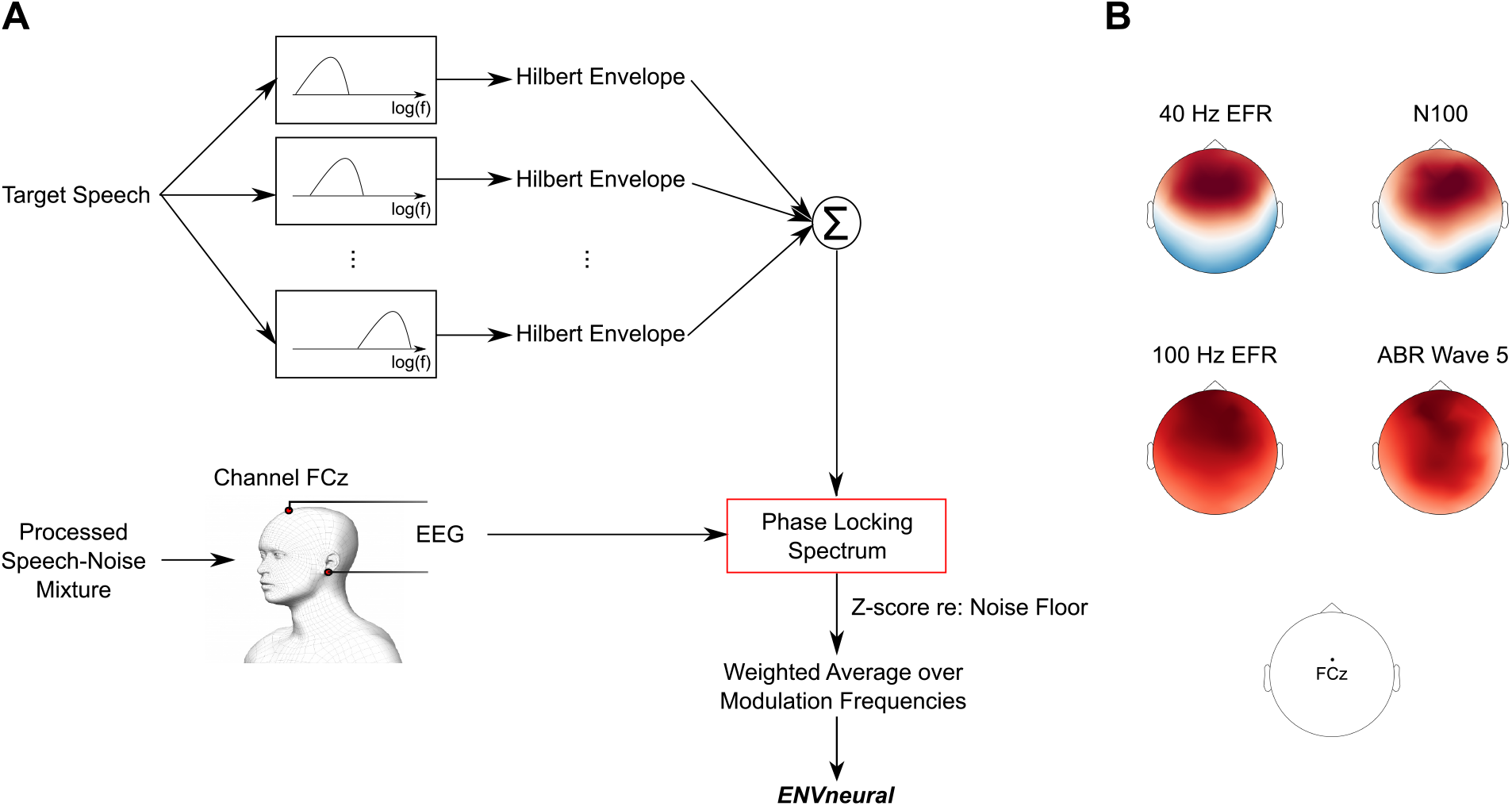
Quantifying the fidelity of target-speech envelope encoding with EEG. **Panel A** illustrates the steps used to quantify target-envelope coding. Target-speech envelopes were extracted using a bank of ten gammatone filters simulating cochlear processing, with roughly log-spaced center frequencies spanning 100–8500 Hz. The envelope at the output of each filter was extracted using the Hilbert transform, and the results were summed across all filters to obtain one overall temporal envelope for each target speech sentence. The fidelity of neural envelope coding of target speech relative to that of background noise (i.e., neural SNR in the envelope domain) was quantified for each experimental condition by computing the phase-locking spectrum between the EEG response in channel FCz and the target-speech envelope across the different trials of that condition (see Equations 1, 2, 3, and 4). The resulting phase-locking spectra were z-scored with respect to an appropriate null distribution of zero phase locking. To obtain a summary metric of neural envelope coding *ENVneural*, the average z-score over all modulation frequencies was computed by weighting the frequencies to compensate for the 1/f transfer function that is observed in EEG measurements. **Panel B** shows that responses from auditory cortex and brainstem are strongest in EEG channel FCz. Data shown are from three different subjects who did not participate in the main experiment but underwent the same screening protocols as the subjects in the main study. Established paradigms for envelope-following responses (EFRs) and onset-evoked potentials (N100 and ABR wave 5) were used to elicit responses from the auditory cortex and brainstem (Picton, 2010). The scalp maps obtained from these responses were normalized such that the amplitudes across channels within each map add to one. The red and blue colors in a scalp map indicate opposite polarities, and the color saturation indicates the normalized amplitude. These scalp maps were used to select the sensor location (FCz) used for all analyses and results presented in this report.

### 2.6 Quantifying EEG-based target-envelope encoding fidelity

We sought to quantify the fidelity (i.e., SNR) of neural envelope encoding of target speech relative to masker fluctuations for each of the eight experimental conditions. The EEG measured in response to our speech-in-noise stimuli reflects not only the neural responses to the target speech and masking noise, but also unrelated brain activity and other EEG measurement noise. To quantify target-envelope coding, we computed the extent to which the EEG response is phase locked to the target-speech envelope using the phase-locking value measure (PLV; Lachaux et al., 1999). We chose this metric because the PLV is monotonically related to the SNR (approximately linearly in the SNR range of *±*6 dB) in the EEG measurements (Bharadwaj and Shinn-Cunningham, 2014) and consequently also to the neural envelope-domain SNR of the target relative to the masker (as sources of noise other than the masker do not vary between conditions). A high PLV between the target-speech envelope and EEG response indicates a consistent phase relationship between those signals, and a low PLV implies little to no relationship between the two signals. Thus, if the EEG response mostly coded target fluctuations (e.g., in a condition with low background noise levels), then the PLV between the EEG signal and the target envelope would be strong. On the other hand, if the EEG response coded mostly masker fluctuations rather than target fluctuations, the PLV would be small. Thus, the PLV captures the envelope-domain SNR with which target envelopes are internally represented relative to modulations in masking sounds and random noise. Note that envelope coding has often been quantified in the literature using a stimulus reconstruction approach, which estimates a linear filter that approximately reconstructs the input speech envelope from EEG responses (e.g., Ding and Simon, 2012; O’Sullivan et al., 2014). Following reconstruction, the proportion of the actual stimulus envelope that is linearly related to the reconstructed envelope is computed as a metric of envelope coding. One disadvantage with this approach is that the first filter estimation step is ill conditioned and necessitates the use of arbitrary regularization techniques (Wong et al., 2018). Our phase-locking measure bypasses this filter estimation step and instead directly captures the proportion of the EEG power that is linearly related to the input speech envelope.

To derive speech envelopes for use in the PLV computation, we used a bank of 10 gammatone filters that mimic cochlear frequency selectivity (Glasberg and Moore, 1990), with center frequencies spanning 100–8500 Hz. The filters were spaced roughly logarithmically, such that their center frequencies had best places that are spaced uniformly along the length of the cochlea according to an established place-frequency map (Greenwood, 1990). Each of the 700 speech sentences used in our study was processed through this filterbank. The envelope of the output of each filter was extracted using the Hilbert transform; the results were summed across all cochlear bands to obtain one overall temporal envelope for each target speech sentence. Note that the single overall envelope obtained by summing across 10 bands is adequate to characterize envelope coding with EEG since EEG does not offer tonotopically resolved information and our focus was not on tonotopic weightings. This is in contrast to the high-resolution procedure crucial for generating vocoded stimuli, as the envelopes conveyed by the periphery are expected to influence the neural processing of target speech. To extract the EEG response to the speech-in-noise stimulus in each trial, a 2.5-s-long epoch that corresponds to the time window during the trial when the target speech was presented was extracted from the overall EEG response in that trial. Epochs corresponding to a particular experimental condition were then pooled over all subjects who performed the condition, to yield a total of 700 epochs per condition. All EEG epochs for a particular condition were paired with the envelopes of the corresponding target speech sentences, and used to calculate the condition-specific PLV measure. PLV was computed in two different ways using custom code adapted from the MNE-Python toolbox (Gramfort et al., 2014), as described below.

The “long-term” PLV spectrum was estimated for each condition using a multi-taper approach (Zhu et al., 2013). Five tapers were used, which resulted in a frequency resolution of 2.4 Hz. The multi-taper PLV estimate minimizes spectral leakage (i.e., reduces mixing of information between far-away frequencies) for any given spectral resolution and is calculated from the Fourier representations of two signals *X*(*f*) and *Y* (*f*) (representing target-speech envelope and EEG response, respectively) as follows:

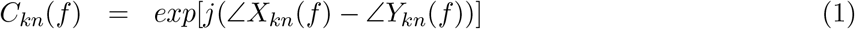

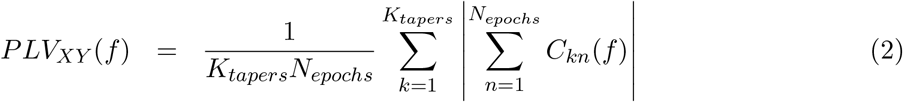

Here, *k* indexes the taper, *n* indexes the epoch, and *f* is modulation frequency.

In addition to the long-term PLV measure described above, we also computed a short-term multi-resolution PLV for modulation frequencies above 7 Hz to account for any modulation masking release that may occur in short time windows. Multi-resolution analyses have been shown to predict intelligibility better than long-term analyses in the case of fluctuating maskers (Jørgensen et al., 2013). A Morlet wavelet was used to compute the EEG and speech spectra in short time windows using seven cycles at each frequency bin (which resulted in a frequency resolution that monotonically decreased with increasing bin center frequency). The window length is inversely proportional to the wavelet center frequency; thus, the number of windows also varied according to frequency (with fewer windows at lower frequencies, and more windows at higher frequencies). Given that each target sentence was a little over 2 s and that each wavelet had seven cycles, the multi-resolution analysis was restricted to 7 Hz and above in order for at least two non-overlapping windows to be resolvable. The multi-resolution PLV is calculated from the Fourier representations of two signals *X*(*f*) and *Y* (*f*) (representing target-speech envelope and EEG response, respectively) as follows:

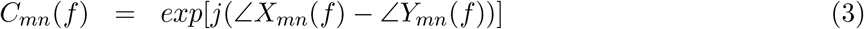

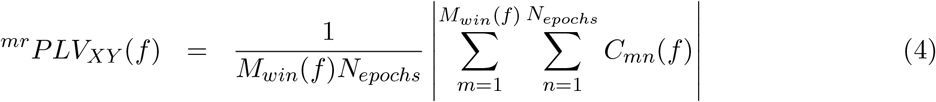

Here, *m* indexes the window, *n* indexes the epoch, and *f* is modulation frequency.

The long-term PLV spectra were averaged within octave-wide modulation bands, spaced half an octave apart. In the case of the multi-resolution PLV computation, we used a similar half-octave spacing when defining the wavelet center frequencies. The binned long-term and multi-resolution PLV spectra thus obtained were z-scored with respect to corresponding null distributions of zero phase locking, which were obtained by pairing EEG trials with mismatching speech trials as described in Section 2.8. The z-scores from the long-term and multi-resolution analyses were thresholded at zero, and then summed at each frequency bin. Then, to obtain a summary metric of neural envelope coding *ENVneural*, the average PLV over all modulation frequency bins was computed after weighting the z-scores in the bins to compensate for the 1/f power transfer function that is characteristic of the SNR of EEG measurements (Stinstra and Peters, 1998; Roß et al., 2000; Buzsáki et al., 2012). Specifically, the z-score in each frequency bin was weighted by a factor proportional to the square root of the bin center frequency, and then the weighted-average z-score across bins was computed. In this way, a separate *ENVneural* metric was quantified for each experimental condition. Note that although different carrier frequency bands and modulation frequencies likely differ in their perceptual importance (Kryter, 1962; Drullman et al., 1994), our *ENVneural* metric does not assign any importance weighting to them. This is because of the possibility that the physiological computations that contribute to our EEG measurements implicitly incorporate such weighting. Figure 2 illustrates the steps used to quantify *ENVneural*.

### 2.7 Testing the hypothesis that the fidelity of target-envelope coding in the brain predicts intelligibility

The hypothesis that the fidelity (i.e., SNR) with which envelopes of target speech are coded in the brain relative to background noise predicts intelligibility (in a quantitative, statistical sense) was tested using a rigorous two-step approach. In the calibration step, a logistic/sigmoid function was used to map the EEG-based *ENVneural* measurements to perceptual intelligibility for speech in stationary noise. This mapping revealed a monotonic relationship between *ENVneural* and intelligibility across the three SiSSN conditions (see Fig. 4B). A crucial test of envelope-based predictions is whether a mapping between *ENVneural* and perceptual intelligibility derived from one type of background noise can be used to estimate intelligibility for novel backgrounds and linear and non-linear distortions applied to the input sounds. In the next step, we predicted intelligibility for speech presented in various novel, realistic backgrounds and distortions from EEG *ENVneural* measurements and by using the mapping created with stationary noise. The following conditions were tested in the prediction step: SiB at 4 dB SNR, SiB at -2 dB SNR, SiB at 6 dB SNR subjected to reverberation, SiB at 4 dB SNR subjected to 64-channel envelope vocoding, and SiB at -6 dB SNR subjected to non-linear denoising (ITFS). Figure 3 illustrates the calibration and prediction steps that were used to test the hypothesis.

**Figure 3.**
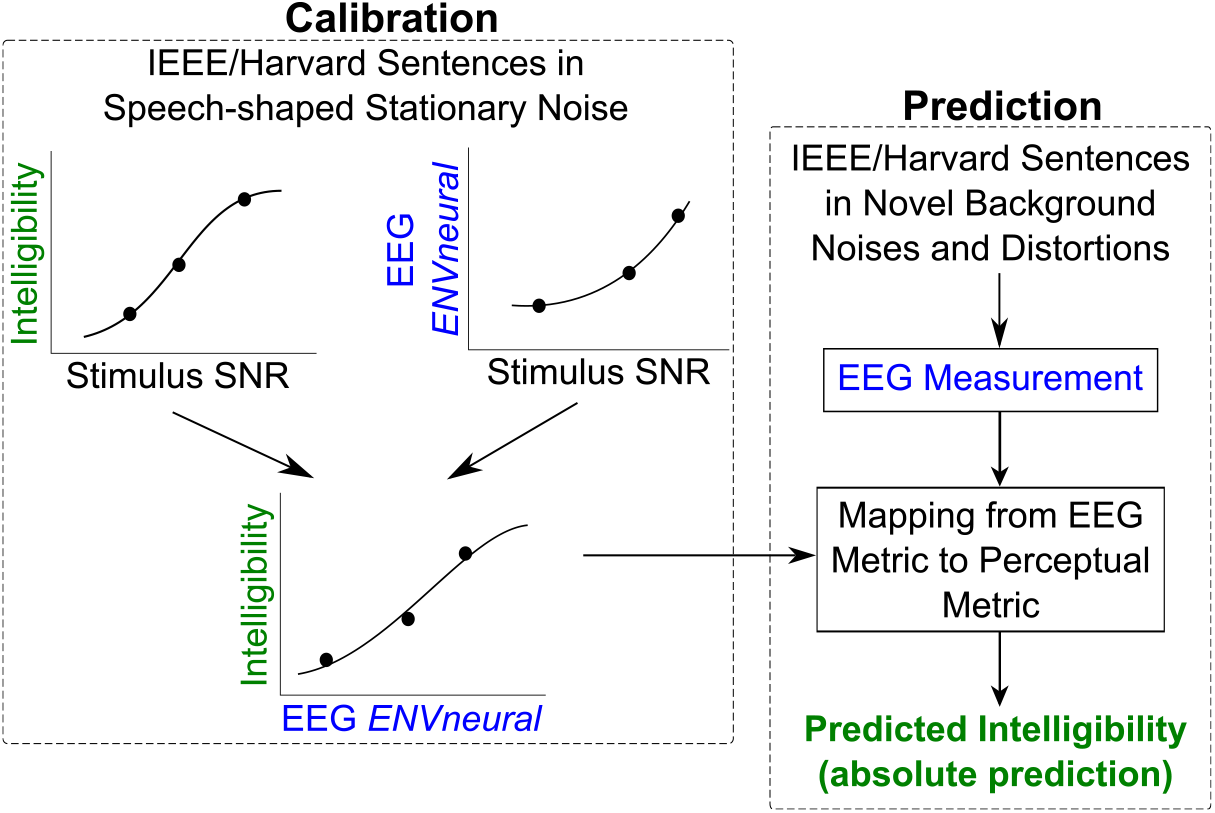
Our rigorous two-step approach to test the hypothesis that the fidelity of neural envelope coding of target speech relative to background noise predicts speech intelligibility (in a quantitative, statistical sense). The first step is a calibration step, where a logistic/sigmoid function was used to map an EEG-based target envelope-coding metric, *ENVneural*, to perceptual intelligibility for speech in stationary noise. In the second step, we used this mapping to blindly predict speech intelligibility in various completely novel realistic background noises and distortions only from EEG-based *ENVneural* measurements.

**Figure 4.**
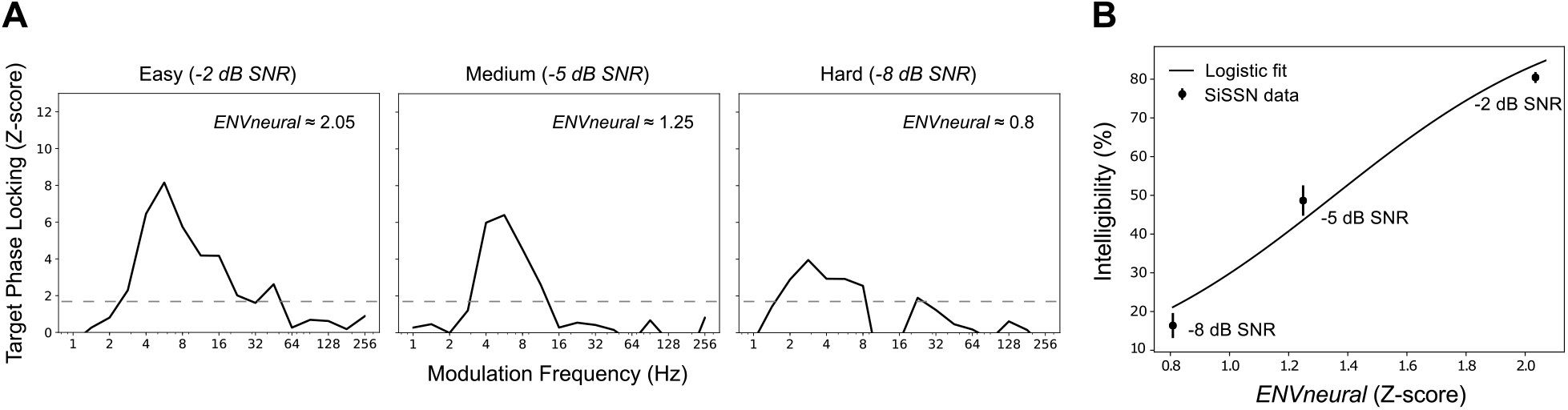
The calibration step: Stationary noise was used to create a mapping between our EEG-based target envelope-coding metric *ENVneural* and perceptual intelligibility. Shown are results from the calibration step of our two-step approach to test the hypothesis that speech intelligibility can be predicted from the fidelity (i.e., SNR) with which target-speech envelopes are coded in the brain relative to background noise. Target-envelope coding fidelity was estimated by computing the phase-locking (PLV) spectrum between the EEG response and the target-speech envelope. **Panel A** shows target PLV spectra (z-scored with respect to a null distribution that is common across conditions) for three SNRs of speech in speech-shaped stationary noise (SiSSN). The dashed lines indicate z = 1.64, i.e., the 95th percentile of the noise floor distribution. Neural envelope coding of the target monotonically decreases with increasing noise (compare the areas under the PLV spectra for -2, -5, and -8 dB SNR). A summary metric of target-envelope coding (i.e., *ENVneural*) was derived separately for each condition by pooling the PLV across modulation frequencies. **Panel B** shows *ENVneural* versus intelligibility measurements (mean and standard error across subjects). The monotonic relationship between *ENVneural* and measured intelligibility across the three SNRs of SiSSN allowed us to fit a sigmoid/logistic function mapping *ENVneural* to intelligibility, as shown, which can then be used for predicting intelligibility from measured *ENVneural* for novel conditions.

### 2.8 Statistical analysis

The distributions for the PLV metric (one for the long-term analysis and another separately for the multi-resolution approach) under the null hypothesis of zero phase locking were obtained using a non-parametric shuffling procedure (Le Van Quyen et al., 2001). Each realization from either null distribution was obtained by following the same computations used to obtain the actual PLV measures, but by pairing EEG response epochs randomly with mismatching speech epochs. That is, when computing the PLV between the speech signal and the EEG signal, the order of epochs for one of the two signals was randomly permuted. This procedure was repeated with 16 distinct randomizations for each experimental condition. Samples were pooled across the 16 randomizations and across all eight conditions to yield a total of 128 realizations from each null distribution. This procedure ensured that the data used in the computation of the null distributions had the same statistical properties as the original speech and EEG signals.

To test the hypothesis that the fidelity of neural envelope coding of target speech relative to that of background noise (i.e., neural SNR in the envelope domain) predicts intelligibility, we computed the Pearson correlation between our EEG-based intelligibility predictions and the actual intelligibility measurements. The p-value for the correlation was derived using Fisher’s approximation (Fisher, 1921).

The noise floor parameters used for computing the z-scores shown in Figure 8 were derived as described in Viswanathan et al. (2019).

### 2.9 Software accessibility

Stimulus presentation was controlled using custom MATLAB (The MathWorks, Inc., Natick, MA) routines. EEG data preprocessing was performed using the open-source software tools MNE-Python (Gramfort et al., 2014) and SNAPsoftware (Bharadwaj, 2018). All further analyses were performed using custom software in Python (Python Software Foundation, Wilmington, DE) and MATLAB. Copies of all custom code can be obtained from the authors.

## 3 Results

### 3.1 Neural envelope-domain SNR in target encoding predicts speech intelligibility over a variety of realistic listening conditions novel to the predictive model

Figure 4 shows results from the calibration step of our two-step approach to test the hypothesis that speech intelligibility can be predicted from the fidelity (i.e., SNR) with which target-speech envelopes are encoded in the brain relative to background noise. As described in Section 2, the fidelity of target-envelope coding in the brain was estimated from the phase-locking spectrum (computed using the phase-locking value; PLV) between the EEG response and the target-speech envelope. Panel A shows target phase-locking (PLV) spectra for three speech in speech-shaped stationary noise (SiSSN) conditions, which correspond to different acoustic SNRs. Comparing the areas under the PLV spectra for -2, -5, and -8 dB SNR shows that the strength of neural envelope coding of the target monotonically decreases with increasing noise. A summary metric of target-envelope coding, *ENVneural*, was derived separately for each condition by pooling the PLV across modulation frequencies (see Section 2). Panel B illustrates the monotonic relationship between *ENVneural* and perceptual intelligibility measurements across the different SNRs of SiSSN. We fit this relationship with a sigmoid/logistic function (shown in the figure) to map *ENVneural* to perceptual intelligibility.

The mapping created in the calibration step was used to predict intelligibility for speech in novel realistic background noises and with different distortions (i.e., conditions not used in calibration), purely from EEG measurements. Figure 5 compares predictions to measured intelligibility for the novel conditions. A total of five novel conditions were tested: speech in four-talker babble (SiB) at SNRs of 4 and -2 dB, SiB at 6 dB SNR subjected to reverberation, SiB at 4 dB SNR subjected to 64-channel envelope vocoding, and SiB at -6 dB SNR subjected to non-linear denoising (using ideal time-frequency segregation; ITFS). Predictions match measured performance closely (*R*^2^ = 0.93, *p* = 0.004), suggesting that envelope coding of the target (relative to the background) in the central auditory system predicts intelligibility. Note that the measurement noise (i.e., background EEG activity unrelated to the target or masker) would be constant across our comparisons. Hence, the variation in *ENVneural* across conditions should primarily reflect the fidelity of target-envelope coding relative to the masker’s internal representation (i.e., the neural modulation-domain SNR). In light of this, the finding that *ENVneural* predicts intelligibility across a range of novel realistic conditions provides neurophysiological evidence for perceptual modulation masking.

**Figure 5.**
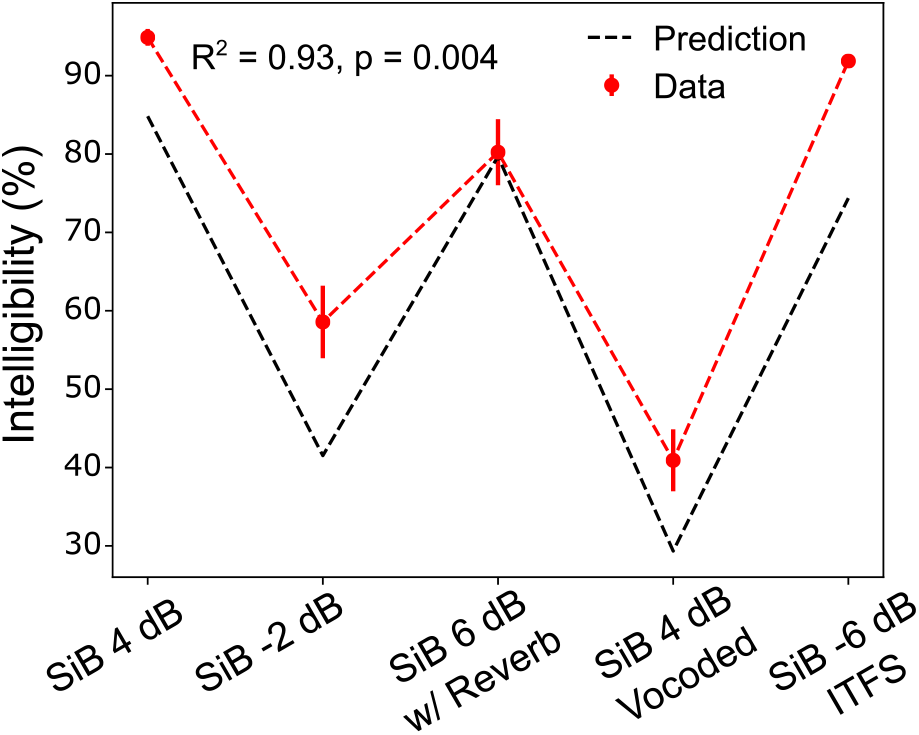
EEG-based target-envelope coding fidelity predicts intelligibility for a variety of realistic conditions not used in calibration. The mapping created using stationary noise (Fig. 4B) was used to predict intelligibility for speech in completely novel realistic background noises and with various distortions, purely from EEG measurements. A total of five novel conditions were tested: speech in four-talker babble (SiB) at SNRs of 4 and -2 dB, SiB at 6 dB SNR subjected to reverberation, SiB at 4 dB SNR subjected to 64-channel envelope vocoding, and SiB at -6 dB SNR subjected to non-linear denoising (using ITFS). Shown are our intelligibility predictions versus actual measurements (mean and standard error across subjects) for these conditions. Predictions match measured performance closely (*R*^2^ = 0.93, *p* = 0.004), suggesting that neural envelope coding of target speech (relative to the background) in the central auditory system predicts intelligibility. Since the measurement noise (i.e., background EEG activity unrelated to the target or masker) would be constant across our comparisons, the variation in *ENVneural* across conditions should primarily reflect the fidelity of target-envelope coding relative to the masker’s internal representation (i.e., the neural modulation-domain SNR). In light of this, the result shown provides neurophysiological evidence for perceptual modulation masking.

### 3.2 The modulation frequencies that contribute to the overall *ENVneural* metric, which predicts intelligibility, depend strongly on the envelope spectrum of the masker

Figure 6 shows target phase-locking (PLV) spectra for two distinct listening conditions: SiSSN, and SiB, as well as modulation spectra for the speech-shaped stationary noise and four-talker babble maskers. The modulation spectra for the maskers were generated by computing the multi-tapered spectral estimates (with five tapers, resulting in a frequency resolution of 2.4 Hz, and 72 trials) of the 2.5-s-long temporal envelope (summed across cochlear bands) of those maskers. Note that the procedure used to generate the masker envelopes was the same as that used to obtain target-speech envelopes for the PLV computation (see Section 2). Comparing the modulation spectrum of speech-shaped stationary noise to the target PLV spectrum for the -2 dB SNR SiSSN condition, we find that speech-shaped stationary noise degrades the representation of high-frequency target modulations more (and low-frequency modulations less), in line with the fact that there is greater power for high-frequency than for low-frequency modulation in stationary noise. On the other hand, comparing the modulation spectrum of four-talker babble to the target PLV spectrum for the 4 dB SNR SiB condition, we see that four-talker babble degrades the representation of low-frequency target modulations more (and high-frequency modulations less). This is consistent with the fact that there is greater power for low-frequency rather than high-frequency modulation in babble. These results show that the spectral profile of EEG-based target-envelope coding fidelity (i.e., the neural envelope-domain SNR in target-speech encoding) is shaped by the masker’s modulation spectrum. This result, in combination with our finding that EEG-based target-envelope coding predicts intelligibility, provides further neurophysiological evidence for perceptual modulation masking. These results also suggest that the modulation frequencies that contribute most to speech intelligibility in realistic listening conditions could lie anywhere in the full continuum from slow prosodic fluctuations to fast pitch-range fluctuations. Previous studies that examined electrophysiological responses to speech in background noise, and how those relate to speech perception, focused on either the cortical tracking of low-frequency envelopes (Ding and Simon, 2014), or on the subcortical tracking of envelope fluctuations in the pitch range (Bidelman, 2017; Shinn-Cunningham et al., 2017). Our findings thus suggest that the prominent use of stationary noise in the previous cortical speech-tracking literature may have been a contributing factor to their focus on low-frequency speech envelopes, i.e., in the so-called “delta” and “theta” ranges.

**Figure 6.**
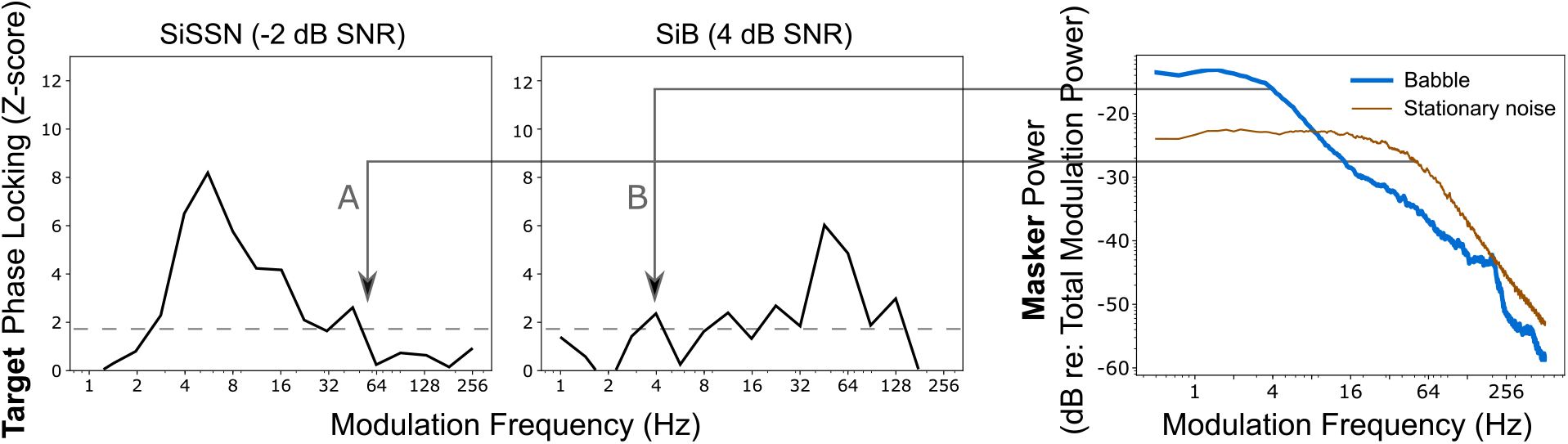
The modulation frequencies that contribute to the overall *ENVneural* metric, which predicts intelligibility, depend strongly on the envelope spectrum of the masker. The target phase-locking (PLV) spectra shown are z-scored with respect to a null distribution that is common across conditions. The dashed lines indicate z = 1.64, i.e., the 95th percentile of the noise floor distribution. Comparing the modulation spectrum of speech-shaped stationary noise (rightmost panel) to the target PLV spectrum for the -2 dB SNR SiSSN condition (A), we find that speech-shaped stationary noise degrades the representation of high-frequency target modulations more (and low-frequency modulations less), in line with stationary noise containing relatively more high-frequency modulation power. In contrast, comparing the modulation spectrum of four-talker babble (rightmost panel) to the target PLV spectrum for the 4 dB SNR SiB condition (B), we show that four-talker babble degrades the representation of low-frequency target modulations more (and high-frequency modulations less), consistent with babble containing relatively more low-frequency modulation power. These results show that the spectral profile of EEG-based target-envelope coding fidelity (i.e., the neural envelope-domain SNR in target-speech encoding) is shaped by the masker’s modulation spectrum. This result, in combination with our finding that EEG-based target-envelope coding predicts intelligibility, provides further neurophysiological evidence for perceptual modulation masking. These results also suggest that the modulation frequencies that contribute most to speech intelligibility in everyday listening could lie anywhere in the full continuum from slow prosodic fluctuations to fast pitch-range fluctuations.

### 3.3 EEG-based envelope coding fidelity and intelligibility are shaped not just by peripheral envelopes but also by TFS

Comparing the SiB at 4 dB SNR (intact) condition with the 64-channel envelope-vocoded SiB at 4 dB SNR in Figure 7, we find that intelligibility and target-envelope coding fidelity in central auditory neurons are both significantly degraded in the vocoded condition. Note, however, that the envelopes at the cochlear level are very similar before and after vocoding (see Section 2) due to the relatively large number of channels (i.e., 64) used in the vocoding process. Despite this, intelligibility is far worse for the vocoded condition compared to the intact condition, demonstrating that the integrity of peripheral envelope cues alone cannot account for speech intelligibility. Importantly, the neural representation of the target envelope in these conditions mirrors these behavioral differences. Thus, the central representation of target envelopes is shaped by factors other than just peripheral envelopes, such as fine-structure-aided segregation mechanisms and selective-attention mechanisms that operate on the segregated representations of target and masker. For example, perceptual cues such as pitch and timbre can aid segregation and selective attention (Darwin, 1997; Micheyl and Oxenham, 2010; Shinn-Cunningham, 2008), but these attributes rely upon stimulus TFS (Smith et al., 2002). When segregation cues are ambiguous, selective attention is impaired, as demonstrated by experiments that engineered conflicting cues (Shinn-Cunningham, 2008; Bressler et al., 2014). The notion that fine-structure cues work together with envelopes in facilitating segregation is consistent with previous psychophysical studies showing that broadband stimuli produce greater pitch-based masking release compared to low-pass or high-pass speech (Oxenham and Simonson, 2009).

**Figure 7.**
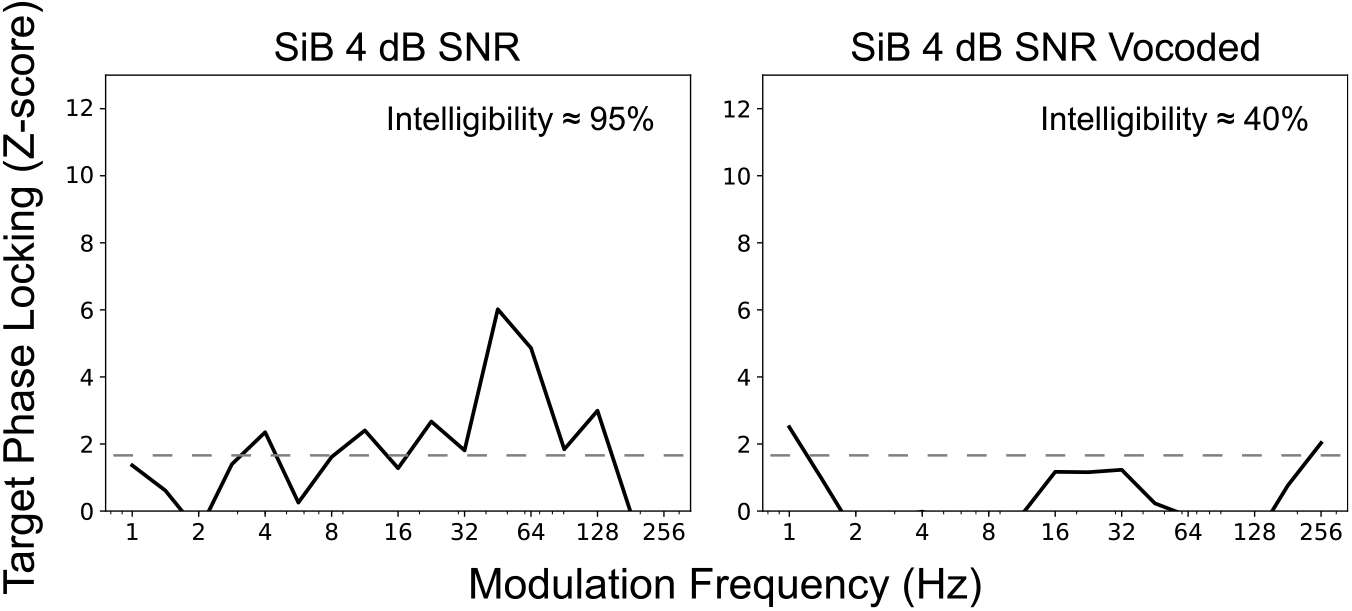
EEG-based envelope coding fidelity and intelligibility are shaped not just by peripheral envelopes, but also by TFS. Comparing the target phase-locking (PLV) spectra (z-scored with respect to a null distribution that is common across conditions) for intact and vocoded SiB at 4 dB SNR shows that 64-channel envelope vocoding significantly degrades envelope coding of the target relative to the background in central auditory neurons, even though the envelopes at the cochlear level are very similar before and after vocoding. Concomitantly, intelligibility is far worse for the vocoded condition compared to the intact condition, demonstrating that the integrity of peripheral envelope cues alone cannot account for speech intelligibility. This result shows that central neural envelope coding and intelligibility are shaped by factors other than just peripheral envelopes, such as stimulus TFS, which supports source segregation and selective attention. Note that the dashed lines indicate z = 1.64, i.e., the 95th percentile of the noise floor distribution.

Many previous studies show that attentional focus, manipulated through subject instruction, can alter central neural envelope coding (e.g., Ding and Simon, 2012; O’Sullivan et al., 2014). Figure 8 illustrates this for a previous study from our lab (reanalysis of data from Viswanathan et al., 2019). Phase locking (averaged over 10 cochlear bands with center frequencies spanning 100–8500 Hz) between the input speech envelope and EEG response depends directly on what speech a listener attends. For the same input speech stream, the speech envelope of a stream is represented more strongly in the brain when that speech is attended to rather than when it is ignored. Thus, central neural envelope coding is shaped by not just peripheral envelopes, but also fine-structure-dependent segregation and selective-attention effects. However, no model of speech intelligibility accounts for this fine-structure contribution.

**Figure 8.**
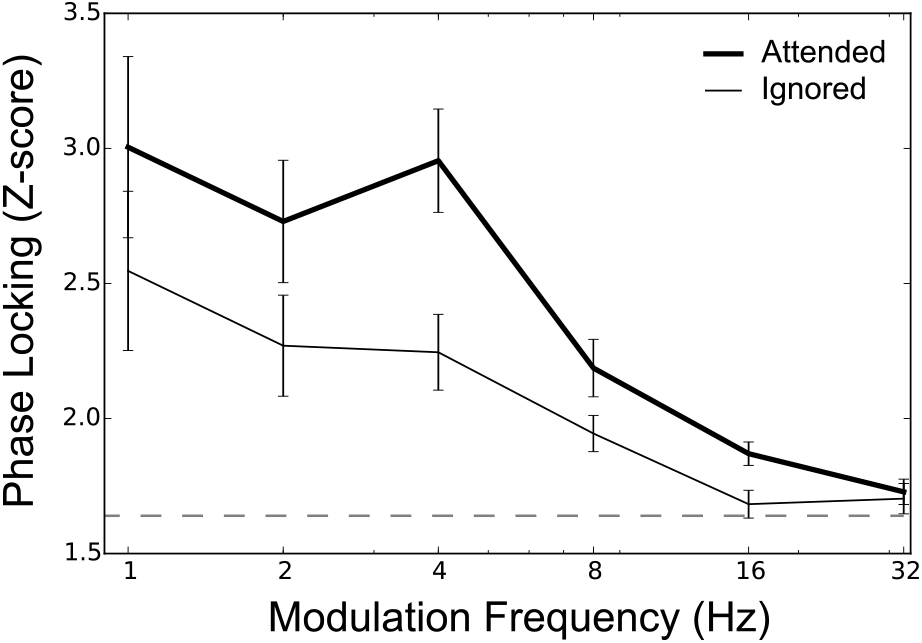
For the same input speech stream, attentional manipulations (via experimental design) alter central neural envelope coding (data reanalyzed from Viswanathan et al., 2019). Subjects were presented with a mixture of two running speech streams, one to be attended to and the other ignored. Selective-attention-dependent phase locking was computed between the input speech envelope and EEG response and averaged over ten cochlear bands with center frequencies spanning 100–8500 Hz. The data shown are the mean and standard errors of phase locking across ten subjects. The dashed line indicates z = 1.64, i.e., the 95th percentile of the noise floor distribution. Speech envelopes are represented more strongly in the brain when speech is attended to versus when the same speech is ignored.

### 3.4 Results support an integrative conceptual model of speech intelligibility

To summarize, our results show that the strength of neural tracking of the target envelope relative to that of the background provides a neural correlate of perceptual interference from a competing sound. Specifically, the ultimate strength of the central auditory system’s encoding of the envelope of target speech relative to other interfering sounds predicts speech intelligibility in a variety of real-world listening conditions. Moreover, we find that the modulation frequencies that contribute to our overall *ENVneural* metric, which predicts intelligibility, depend strongly on the envelope spectrum of the masker and the scene acoustics. Note that modulation-frequency-specific effects can arise from within-channel masking where the masker contains elements that share the same carrier and modulation frequency as some target elements (Jørgensen and Dau, 2011) or from cross-channel interference where masker elements that are coherently modulated with target elements interfere with target coding and perception (Apoux and Bacon, 2008). Our EEG-based metric does not distinguish between these distinct forms of temporal-coherence-based effects. Rather, our results provide evidence that some combination of the two shapes scene analysis and speech perception in noise. Our results also provide direct neural evidence that TFS cues affect how well neural responses in the central auditory system encode the envelope of target speech, likely by aiding in successful source segregation (Darwin, 1997; Oxenham and Simonson, 2009; Micheyl and Oxenham, 2010) and selective attention (which can operate on the internal representation of segregated target and masker objects to boost the neural representation of the target relative to the masker; Ding and Simon, 2012; O’Sullivan et al., 2014; Viswanathan et al., 2019). Taken together, our neurophysiological results support the theory that scene analysis and attentive selection of target speech are influenced by both modulation masking and TFS, consistent with the broader temporal coherence theory. These ideas motivate our conceptual model of speech intelligibility (Fig. 9), which consolidates these elements into a single framework.

**Figure 9.**
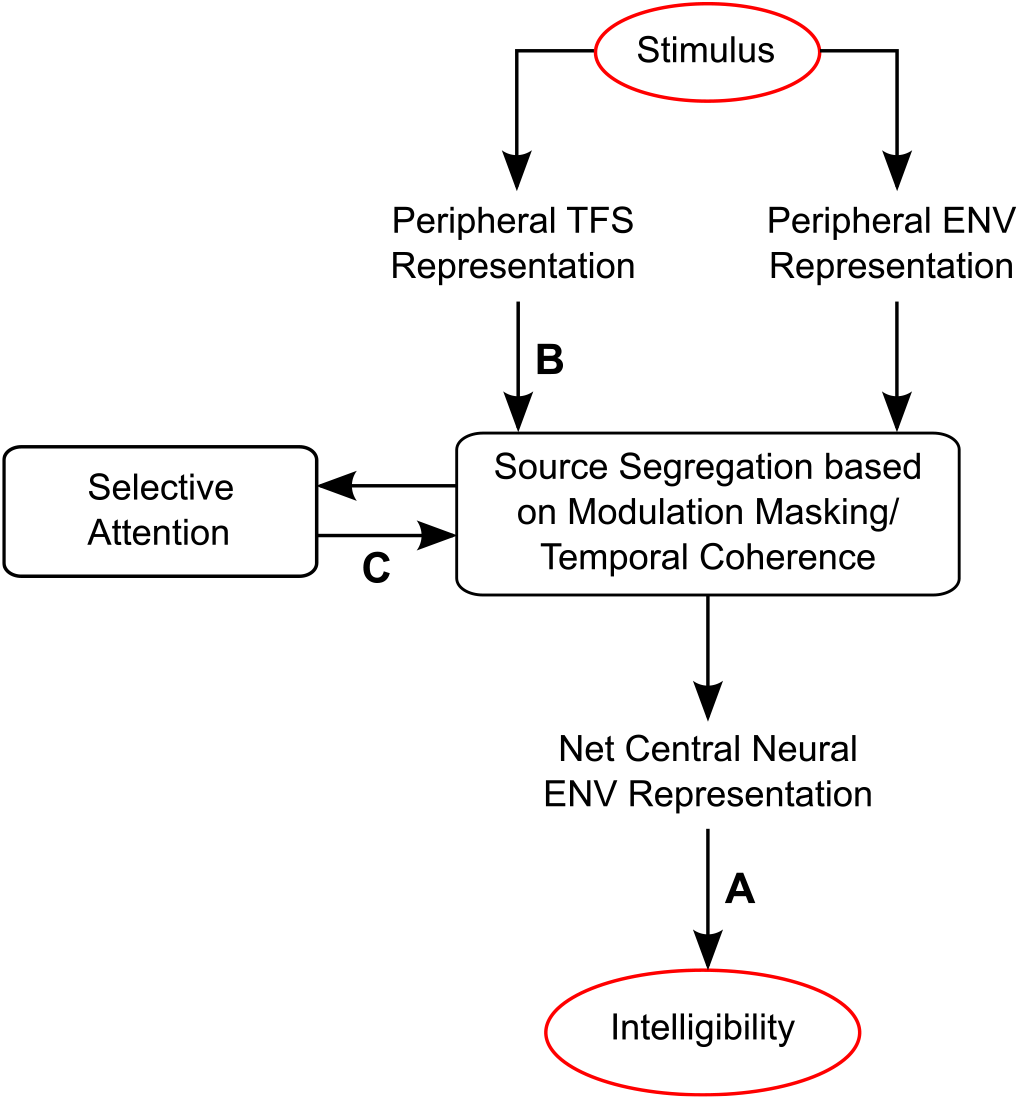
Results support an integrative conceptual model of speech intelligibility. Taken together, our results support this integrative conceptual model of speech intelligibility in that they clarify what internal representation is predictive of speech intelligibility and how that representation is related to the acoustics of the auditory scene and cognitive variables. Our results show that the strength of the net envelope (ENV) coding of target speech relative to other interfering sounds in the central auditory system predicts intelligibility in a variety of real-world listening conditions (arrow A). The modulation frequencies that contribute to these EEG-based intelligibility predictions depend strongly on the envelope spectrum of the masker and the scene acoustics. TFS cues (arrow B) also affect how well neural responses in the central auditory system encode the envelope of target speech, likely by aiding in source segregation (Darwin, 1997; Oxenham and Simonson, 2009; Micheyl and Oxenham, 2010). Selective attention can then operate effectively on the distinct representations of segregated target and masker objects (arrow C) to boost the neural representation of the target relative to the masker (Ding and Simon, 2012; O’Sullivan et al., 2014; Viswanathan et al., 2019). Taken together, our results support the theory that scene analysis and attentive selection of target speech are influenced by both modulation masking and TFS, consistent with the broader temporal coherence theory.

## 4 Discussion

The present study systematically examined how neural encoding of target speech in the central auditory system varied as characteristics of the scene acoustics and background noise were manipulated, and how these neural metrics are related to speech intelligibility. Our results provide support for the temporal coherence theory of scene analysis (Elhilali et al., 2009) in that (i) our EEG-based target-envelope coding metric, which predicts intelligibility, is strongly influenced by the envelopes in background noise in a modulation-frequency-specific manner, and (ii) the availability of intact TFS enhances target-envelope coding.

A key result here is that the neural envelope-domain SNR in target encoding predicts intelligibility (in a quantitative, statistical sense) for a range of strategically chosen real-world conditions that are completely novel to the prediction model. Furthermore, the set of target-envelope frequencies that contribute to our EEG-based intelligibility prediction depends strongly on the envelope frequencies contained in the background sounds. These results together suggest that modulation masking may be fundamentally important for speech perception in noise, thus validating previous behavioral studies (Bacon and Grantham, 1989; Stone and Moore, 2014) and current speech-intelligibility models (Dubbelboer and Houtgast, 2008; Relaño-Iborra et al., 2016) with neurophysiological evidence. Note, however, that our results do not directly provide evidence of neural modulation filter banks (Jørgensen et al., 2013; Relaño-Iborra et al., 2016). Another mechanism by which modulation masking could occur is through temporal-coherence-based binding across a distributed assembly of neurons (Eckhorn et al., 1990; Eggermont, 2006). Through this mechanism, those envelope and fine-structure frequencies of the target that are temporally coherent with components of the masker may get bound together (i.e., a failure of source segregation), which in turn can lead to degraded target representation and perceptual modulation masking at those specific frequencies. Indeed, there is evidence that the redundancy in temporal pitch information across low-frequency resolved harmonics and high-frequency envelopes is more effective in facilitating masking release than what is obtained from either of them individually (Oxenham and Simonson, 2009). Our findings underscore the need for further research into the neural circuit-level computations that support such complex integration of various temporal cues during active listening.

Previous psychophysical studies with carefully processed speech stimuli in quiet (Shannon et al., 1995; Smith et al., 2002; Elliott and Theunissen, 2009) and the success of envelope-based cochlear implants in quiet backgrounds (Wilson and Dorman, 2008) suggest that envelope coding is fundamental for speech perception. However, a more general and rigorous test of this hypothesis requires an examination of whether or not envelope coding predicts intelligibility for the average listener over a range of realistic listening conditions not used by the predictive model. Some prior studies compared individual variations in envelope coding to intelligibility; these used just one type of masker, such as stationary noise (e.g., Ding and Simon, 2013; Vanthornhout et al., 2018) or a multi-talker interferer (e.g., Bharadwaj et al., 2015). In contrast, we were able to predict intelligibility in a variety of novel ecologically relevant conditions from just average neural metrics, learning the prediction model from the independent stationary-noise condition. Despite EEG measurement noise or any errors introduced due to variability in intelligibility measurements in the calibration step, our EEG-based predictions closely track (*R*^2^ = 0.93, *p* = 0.004) the overall pattern in measured intelligibility across conditions (Fig. 5C). This is in fact stronger evidence that neural envelope coding is a correlate of speech intelligibility than being able to correlate individual differences in neural coding and behavior, both because pooling across subjects (who differ in performance) adds noise to the metrics we computed, and more importantly because correlated individual differences in EEG and behavior could easily arise from extraneous factors such as motivation, attention, level of arousal, etc. that are unrelated to envelope coding (Bharadwaj et al., 2019).

Another fundamental insight from the present study is that central neural envelope coding depends not only on envelopes conveyed by the inner ear, but also on the TFS. Although this result was reported by Ding et al. (2014), they used four- and eight-channel envelope vocoding to degrade the TFS; this broadband vocoding is in contrast to the high-resolution (64-channel) envelope vocoding that we used here. As demonstrated in Section 2, low-resolution vocoding introduces spurious envelopes (not present in the original stimuli) during cochlear filtering of the noise carrier used in vocoding (Gilbert and Lorenzi, 2006). These spurious envelopes introduced within individual frequency channels are large enough to degrade neural envelope coding in a way that is easily perceptible (Swaminathan and Heinz, 2012), and could account for the reduced cortical target-envelope coding they observed (Fig. 1). Previous behavioral work (Dorman et al., 1998; Qin and Oxenham, 2003) also shows that increasing the number of noise-vocoding channels beyond eight considerably improves speech intelligibility in noise, despite the fact that the TFS is uninformative regardless of the number of channels used in vocoding. Together, these results demonstrate that it is necessary to use high-resolution vocoding, as we do here, to unambiguously attribute effects to TFS cues rather than spurious envelopes. Our 64-channel vocoding procedure leaves place coding and cochlear-level envelopes largely intact (Fig. 1), not only at filters with center frequencies matching the vocoder sub-bands, but also at filters that are midway between adjacent sub-bands. Thus, it is unlikely that peripheral envelope distortion can account for degraded central neural envelope coding and intelligibility in the present study. These neurophysiological results are consistent with previous behavioral studies showing that fine-structure cues aid in scene segregation and selective processing of target speech (Darwin, 1997; Shinn-Cunningham, 2008; Oxenham and Simonson, 2009). The present study also points to a need for more sophisticated speech-intelligibility models that account for the various scene-analysis mechanisms in play to better predict performance across a wider range of conditions (including vocoded speech-in-noise; Steinmetzger et al., 2019).

Our EEG-based two-step approach can be used to test and refine speech-intelligibility models. A major strength of this approach is that it intrinsically factors in listener attributes (e.g., hearing-loss profile, language experience) and listening state (e.g., focus of attention) in addition to purely stimulus-dependent aspects of coding. How different factors contribute to speech perception can be systematically investigated by characterizing how much each factor contributes to the neural response and how the respective contributions are weighted to best predict intelligibility across various conditions. For instance, here we studied how an acoustic aspect of the stimulus (temporal envelope) is coded in the central auditory system by deriving EEG metrics from scalp locations that strongly reflect auditory cortex and brainstem contributions. In addition, higher-order stimulus features, such as phonemic (categorical) processing (Di Liberto et al., 2018a) and semantic composition (Brodbeck et al., 2018) may be studied in future experiments, perhaps by extending our analyses to other brain regions (Du et al., 2014; Di Liberto et al., 2018b; Kim et al., 2020). Similarly, by studying individuals with different peripheral pathophysiologies, the effects of various forms of hearing loss on neural coding and intelligibility can also be characterized (Swaminathan and Heinz, 2011; Rallapalli and Heinz, 2016).

One limitation of our approach is that stimulus-related responses in the EEG can be captured and separated from background brain activity only by virtue of their temporal signature. If certain features are encoded through abstract rate-based representations or through different activation profiles within a spatially distributed organization of receptive fields, our approach cannot readily account for them. For example, cortical neurons represent temporal envelopes not only through phase locking, but also through rate-based tuning (Wang et al., 2008). Furthermore, place/spectral cues are important for speech recognition (Boothroyd et al., 1996; Shannon et al., 1998; Elhilali et al., 2003), whereas EEG measurements are not place specific but instead reflect population neural activity. One consequence of this fact is that our metrics cannot distinguish between within-channel modulation masking where the masker contains elements that share the same carrier and modulation frequency as some target elements (Jørgensen and Dau, 2011) and cross-channel modulation interference where masker elements that are coherently modulated with target elements interfere with target coding and perception (Apoux and Bacon, 2008). Future EEG studies should attempt to delineate cross-channel versus within-channel effects in scene analysis and speech perception, perhaps by employing frequency-separated target speech and masking sounds. Despite these issues, we find that neural encoding of temporal envelopes can account for much of the intelligibility variations seen across the stimulus conditions tested in this study. This may be because (i) although EEG signals cannot be readily used to decode the perceived phonemes, they can adequately capture the overall fidelity with which envelopes are coded despite the lack of tonotopic specificity, and (ii) at slow modulation frequencies, temporal coding may be a prominent mechanism in the cortex (Wang et al., 2008), and at faster modulation frequencies (e.g., in the pitch range), our metric also includes a small contribution from subcortical portions of the auditory pathway where the coding of envelopes is largely temporal (Joris et al., 2004).

## 5 Conclusion

By combining human EEG with simultaneous speech intelligibility measurements, we find that the neural representation of target-speech envelopes is shaped by masker modulations and that this net target-to-masker envelope-domain SNR in central auditory neurons predicts intelligibility over a variety of ecologically relevant conditions novel to the predictive model. Moreover, TFS cues can influence this envelope encoding in the brain, likely by supporting source segregation and selective attention. Finally, a conceptual model of speech intelligibility that integrates these ideas is proposed.

## 6 Acknowledgments

This work was sponsored by grants from the National Institutes of Health [Grant Nos. F31DC017381 (to V.V.), R01DC009838 (to M.G.H.), R01DC015989 (to H.M.B.), and 9605702, R01DC013825 (to B.G.S.-C.)] and from Action on Hearing Loss [Grant No. G72 (to M.G.H.)].

## 7 Author Contributions

V.V., H.M.B., B.G.S.-C., and M.G.H. designed research; V.V. performed research; V.V. analyzed data; V.V. wrote the paper with edits from H.M.B., B.G.S.-C., and M.G.H.

